# Development of a novel glycoengineering platform for the rapid production of conjugate vaccines

**DOI:** 10.1101/2021.11.25.470047

**Authors:** Sherif Abouelhadid, Elizabeth Atkins, Emily Kay, Ian Passmore, Simon J North, Burhan Lehri, Paul Hitchen, Eirik Bakke, Mohammed Rahman, Janine Bosse, Yanwen Li, Vanessa S. Terra, Paul Langford, Anne Dell, Brendan W Wren, Jon Cuccui

**Affiliations:** Department of Infection Biology, London School of Hygiene & Tropical Medicine, WC1E 7HT, UK; Department of Life Sciences, Imperial College London, SW7 2AZ, UK; Department of Infectious Diseases, Imperial College London, W2 1NY, UK

## Abstract

Antimicrobial resistance (AMR) is threatening the lives of millions worldwide. Antibiotics which once saved countless lives, are now failing, ushering in vaccines development as a current global imperative. Conjugate vaccines produced either by chemical synthesis or biologically in *Escherichia coli* cells, have been demonstrated to be safe and efficacious in protection against several deadly bacterial diseases. However, conjugate vaccines assembly and production have several shortcomings which hinders their wider availability. Here, we developed a tool, Mobile-element Assisted Glycoconjugation by Insertion on Chromosome, MAGIC, a novel method that overcomes the limitations of the current conjugate vaccine design method(s). We demonstrate at least 2-fold increase in glycoconjugate yield via MAGIC when compared to conventional bioconjugate method(s). Furthermore, the modularity of the MAGIC platform also allowed us to perform glycoengineering in genetically intractable bacterial species other than *E. coli*. The MAGIC system promises a rapid, robust and versatile method to develop vaccines against bacteria, especially AMR pathogens, and could be applied for biopreparedness.

## Introduction

The alarming rise in antimicrobial resistance necessitates global efforts to prevent a future health crisis. For more than half a century, antibiotics were considered the first line of defence against bacterial pathogens (1). However, the spread of antibiotic resistance amongst pathogenic bacteria entails considerable efforts to look for antibiotic alternatives. Vaccines have been successful in curbing infectious diseases for decades, not only among adults but also among children and the elderly, thus saving millions of lives worldwide (2). According to the Market Information for Access to Vaccines (MI4A), World Health Organization, the vaccines market is estimated to be worth approximately $33 billion in 2019 (3). Current biotechnological platforms however, might not be able to fulfil the vaccines supply demand. To satisfy the market’s demand for conjugate vaccines, to protect humanity from a foreseeable pandemic, and to be able to tailor novel efficacious vaccines at lower cost, significant biotechnological innovation is needed.

Glycoconjugate vaccines are considered to be one of the safest and most effective tools to combat serious infectious diseases including bacterial meningitis and pneumonia (4). Conjugation is achieved by linking glycans (carbohydrate moiety), either chemically or enzymatically, to proteins via covalent bonds. This leads to a T-cell dependent immune response, offering excellent protection in people of all ages (5). Traditionally chemical approaches to produce glycoconjugate vaccine involve the activation of functional groups on the glycan and protein that are linked chemically in a multi-step method that is expensive and laborious, requiring several rounds of purification after each step (6). Additionally, chemical conjugation methods such as reductive amination can alter the polysaccharide epitope, affecting the immunogenicity of the glycoconjugate against the disease, besides its inherent batch-to-batch variation (7). Biological conjugation (bioconjugation) offers an excellent alternative to chemical conjugation. It is based on using a bacterial cell, usually *E. coli*, as a chassis to express a pathway that encodes the desired bacterial polysaccharide, carrier protein, and an oligosaccharytransferase enzyme, OST, that catalyses the conjugation process(6).

The advent of the bacterial bioconjugation method allowed several protein glycan vaccine combinations to be successfully developed, emphasizing its immense potential to become the preferred method to develop glycoconjugate vaccines in the future (4,8–12). However, several challenges remain. Firstly, the process places significant metabolic stress on the *E. coli* cell, vaccine micro-factory, due to the expression of orthogonal pathways (13). This process requires the prior genetic and structural information of the polysaccharide structure of choice. Secondly, the use of three independent replicons has limitations due to incompatibility of plasmid origins of replication and antibiotic selection markers which may lead to the plasmid loss that results in reduction in glycoconjugate yield (8,10,14). Thirdly, reports have demonstrated that the expression of the OST PglB, that catalyses the linking of glycans to carrier proteins, has a detrimental effect on bacterial growth, thus decreasing cellular fitness to produce glycoconjugates (13,15). All this together results in a low biomass which often translates to a reduction in the vaccine yield. Consequently, this leads to an increase in the production cost of a glycoconjugate vaccine, making it unaffordable in low-income countries where they are most needed, putting millions of lives at risk as a result of vaccines inequity (6).

Previous attempts to engineer robust glycoengineering host strains using homologous recombination have had limited success (15). Although this technology managed to moderately boost glycoprotein production and reduce the dependence on plasmids, it suffers from major drawbacks. Firstly, the method is slow and requires the successful expression of recombinase systems from plasmids to allow chromosomal integration of glycoengineering components. Secondly, it cannot be applied to other Gram-negative bacteria since prior knowledge of the genome sequence is required to allow for the design of homologous arms for homologous recombination to occur. Thirdly, the scarcity of genetic manipulation tools, which are available for few bacteria, impede the wide use of homologous recombination platforms.

Here, we present a novel platform to overcome some of the limitations of bioconjugation by creating a modular system to rapidly develop conjugate vaccine candidates. This platform could be biologically tailored in a “plug-and-play” manner to allow the integration and stable expression of the glycoengineering component(s) not only in *E. coli* but also in other Gram-negative bacteria. We term this technique, Mobile-element Assisted Glycoconjugation by Insertion on Chromosome (MAGIC). We demonstrate how this platform enables the rapid assembly of stable glycoconjugate combinations. We also report that once a bacterial cell has undergone the MAGIC process, it can be used as a chassis strain to achieve superior glycoconjugate yields and higher glycosylation efficiency when compared to the traditional three plasmid-based bioconjugation methods, and cell free glycosylation method. Furthermore, integration of the OST into host *E. coli* was shown to alleviate much of the metabolic burden from the bacterium that was translated into increase of biomass and glycoconjugate yield. In addition, we demonstrate the versatility and robustness of MAGIC in not only glycoengineering *E. coli* strains but also in other genetically intractable host bacteria such as *Citrobacter* species. As an exemplar, we demonstrate how MAGIC could provide a first line of defence in an *E. coli* O157 outbreak scenario by developing a candidate glycoconjugate vaccine in less than a week. The modular nature of MAGIC highlights its applicability as a tool for biopreparedness especially against emerging multi-drug resistant bacteria.

## RESULTS

### Assessment of MAGIC in developing a conjugate vaccine against *Francisella tularensis*

To demonstrate the proof-of-principle of MAGIC application in glycoconjugate production, we first constructed *E. coli* MAGIC v.1, based on inducible *pglB* under a P_tac_ promoter system (16) Fig 1, A. We applied MAGIC v.1 to all major glycoengineering strains such as, *E. coli* W3110, *E. coli* SDB1, *E. coli* SΦ874, *E. coli* SCM3, *E. coli* SCM6, *E. coli* SCM7, and *E. coli* CLM24. This was achieved by overnight conjugation, and subsequent culturing, under antibiotic selection, to confirm *pglB* integration on the chromosome. The stability of this integration was confirmed by subculturing *E. coli* MAGIC v.1 more than 10 times without any antibiotic selection and demonstrating that the insertion site had remained intact (data not shown). We then tested *E. coli* MAGIC v.1 in developing a vaccine against *Francisella tularensis* SchuS4. The bacterium *F. tularensis* is categorized as a scheduled bioterrorism class A threat agent due to its high infectivity, low infectious dose, and ease of aerosol distribution (17). The O-antigen of *F. tularensis*, designated here as Ft O-Ag, consists of the repeating unit of the tetrasaccharide [2)-β-Qui4NFm-(1→4)-α-GalNAcAN-(1→4)-α-GalNAcAN-(1→3)-β-QuiNAc-(1→]. Previous studies demonstrated that animals vaccinated by either Ft-LPS or Ft O-Ag glycoconjugate were protected against *F. tularensis* infection (10). Initially, we established *E. coli* as a positive control, designated here as *E. coli* gCmeA bioconjugation Ft O-Ag, expressing Ft O-Ag biosynthetic pathway, PglB under P_tac_ promoter from pEXT21, and CmeA as a model carrier protein. The periplasmic accessory protein CmeA glycoprotein from *Campylobacter jejuni* has been long used as a model carrier protein to investigate glycoengineering components. The glycoprotein carries two native glycosylation sites, ^121^DF<u>N</u>RS^125^ and ^271^DN<u>N</u>NS^275^ (18). A C-terminal 6xHis tag was added to CmeA to facilitate its purification via immobilised metal ion chromatography (IMAC) and Western blot analysis. We used a *waaL* deficient *E. coli* strain, CLM24, considered optimal for glycoconjugate production since the O-antigen ligase gene, *waaL*, is deleted, preventing competition with PglB for Und-PP linked substrate (19).

**Figure 1.**
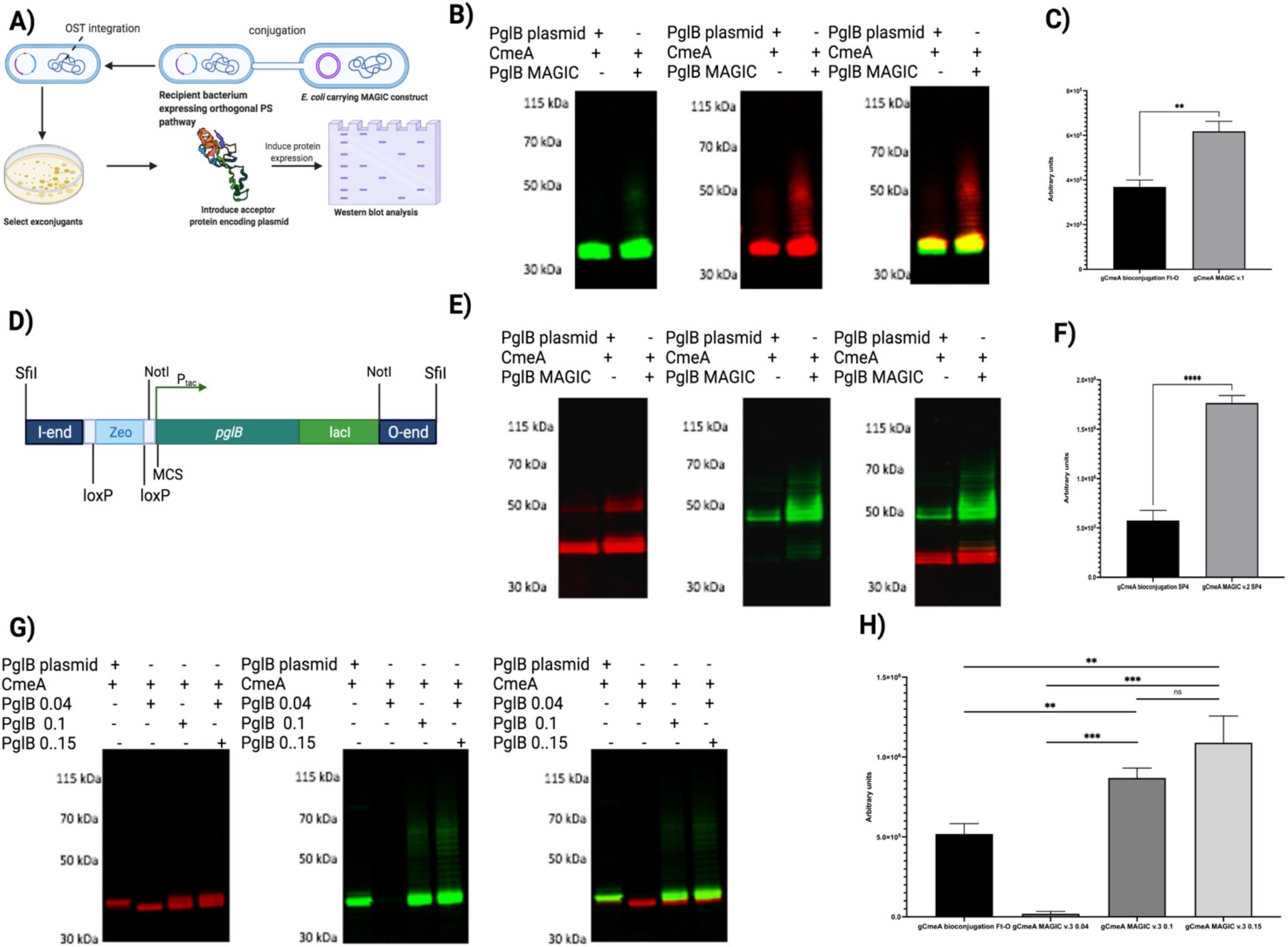
Glycoconjugate production in *E. coli* MAGIC strains compared to conventional bioconjugation method. A, Schematic diagram of developing of constructing *E. coli* MAGIC; B, Western blot of 5 µg His-tagged CmeA protein purified by nickel affinity chromatography. Biological samples were separated on a Bolt 4-12% bis-tris gel (Invitrogen) with MOPS buffer and transferred to nitrocellulose membrane with an iBlot 2 dry blotting system. The membrane was probed with anti-His (Invitrogen) and anti-Ft-O antigen monoclonal antibody (abcam) and detected with fluorescently labelled secondary antisera (green-His, red-Ft-O-antigen) on a LiCor Odyssey scanner.; C, densitometry analysis of glycoconjugate production in *E. coli* MAGIC v.1 Ft-O compared to *E. coli* bioconjugation Ft-O.; D, Design and construction of MAGIC v.2; E, Western blot of 5 µg His-tagged CmeA protein purified by nickel affinity chromatography. Biological samples were separated on a Bolt 4-12% bis-tris gel (Invitrogen) with MOPS buffer and transferred to nitrocellulose membrane with an iBlot 2 dry blotting system. The membrane was probed with anti-His (Invitrogen) and anti-SP4 (Statens) and detected with fluorescently labelled secondary antisera (red-His, green-anti-SP4) on a LiCor Oddysey scanner.; F, densitometry analysis of glycoconjugate production in *E. coli* MAGIC v.2 SP4 compared to *E. coli* bioconjugation SP4.; G, Western blot of 5 µg His-tagged CmeA protein purified by nickel affinity chromatography. Biological samples were separated on a Bolt 4-12% bis-tris gel (Invitrogen) with MOPS buffer and transferred to nitrocellulose membrane with an iBlot 2 dry blotting system. The membrane was probed with anti-His (Invitrogen) and anti-Ft-O antigen monoclonal antibody (abcam) and detected with fluorescently labelled secondary antisera (green-His, red-Ft-O-antigen) on a LiCor Oddysey scanner; H, densitometry analysis of glycoconjugate production in *E. coli* MAGIC v.3 Ft-O compared to *E. coli* bioconjugation Ft-O. Densitometry analysis of glycoconjugate was done in three biological replicates. ns,*p*>0.05; *\*,p*<0.05, **,*p*<0.01, ***, *p*<0.001.

We assembled *E. coli* CLM24 MAGIC v.1, designated here as gCmeA MAGIC v.1 Ft O-Ag as summarized in Fig 1, A. To assess the performance of *E. coli* CLM24 MAGIC v.1 against gCmeA bioconjugation Ft O-Ag. Both *E. coli* CLM24 variants were grown overnight and subcultured the following day. Bioconjugation was initiated by induction of *pglB* expression at OD_600_ of O.4-0.5. Under shake flask culture conditions, we demonstrated that *E. coli* gCmeA bioconjugation Ft O-Ag, carrying the three plasmids system, induced with 1 mM IPTG overnight, reached a maximum OD_600nm_ of 1.2 whilst gCmeA MAGIC v.1 Ft O-Ag grew, on average, to an OD_600_ of 1.7 (average of 41% increase in cell density). Western blot analysis of representative CmeA affinity purified from both *E. coli* strains reacted positively when probed with *F. tularensis* O-antigen antibody. Interestingly, a visible ladder distinctive of *F. tularensis* O-antigen was observed only in gCmeA MAGIC v.1 Ft O-Ag when probed by either anti-his antibody or O-antigen antibody Fig 1, B. Glycoprotein yield was quantified using image densitometry (glycoprotein/glycoprotein + unglycosylated protein *100) from three biological replicates Fig S1. An approximate 2-fold increase in glycoprotein production in gCmeA MAGIC v.1 Ft O-Ag was observed when compared to its counterpart, with glycosylation efficiency increasing from 77.2% ± 14 using traditional three plasmids bioconjugation method to 90.2 % ± 2.9 in gCmeA MAGIC v.1 Ft O-Ag Fig 1, C. Taken together these results show that MAGIC is an effective tool in vaccine production. It also shows that MAGIC alleviates certain cellular burdens caused by the glycoengineering component(s) thus allowing for an increased cell density and higher glycoprotein production.

### Improving of MAGIC v.1

Guided by these results, we sought to improve MAGIC v.1. Firstly, by enhancing the assembly process of MAGIC strains by designing a modular system that is more compatible with loading of the glycoengineering component. Secondly, reducing the dependence of antibiotics as a counter selection marker in the glycoconjugate production process. Thirdly, eliminating any unnecessary DNA sequence that increased the size of MAGIC, making chromosomal integration more efficient. To achieve this, we synthesized DNA having the I and O end of MAGIC v.1 and we reduced the cargo size by selecting a small DNA fragment encoding Zeocin^®^ resistance cassette (359 bp), replacing a larger antibiotic resistant cassette, kanamycin cassette (816 bp). The Zeocin^®^ resistance cassette is flanked by *loxp* site to allow removal of antibiotic selection marker once it is integrated on the chromosome. Secondly, we added unique restriction enzyme sites for facile loading of cargo on a high copy pUC vector before transferring onto the chromosomal delivery construct. We designate this as MAGIC v.2 Fig 1, D.

We sought to test MAGIC v.2 in developing a vaccine against the bacterium *S. pneumoniae*, a leading cause of pneumonia and meningitis worldwide (20) Pneumococcal capsular polysaccharide is one of the main virulence factors and a major component of the vaccine currently in use (PPSV23 and PCV10, 13 and 15). Currently, vaccines against *S. pneumoniae* are produced via chemical conjugation (21). Previously, we demonstrated that a biologically conjugated vaccine against *S. pneumoniae* confers protection and increased survival rate in laboratory animals (14). We sought to apply MAGIC v.2 in enhancing the production of a *S. pneumoniae* vaccine. As a control to this experiment, *E. coli* W3110 CmeA-Sp4 was assembled. This strain expresses the orthogonal pathway of *S. pneumoniae* serotype 4 (Sp4), which consists of Pyr-Glc-(1→3)-α-ManNAc-(1→3)-β-FucNAc-(1→3)-α-GlcNAc (22), PglB under P_tac_ promotor from a single copy plasmid pEXT22, and a carrier protein CmeA. Previous attempts to generate a Sp4-CmeA glycoconjugate in CLM24 and W3110 carrying *pglB* expressed from pEXT21 were unsuccessful. We constructed *E. coli* W3110 MAGIC v.2 as detailed in the methods section and removed the antibiotic resistance cassette (Zeocin^®^) using the *cre/loxP* system. Successive subculturing in the absence of antibiotics did not result in the loss of the OST following chromosomal integration due to MAGIC v.2. Next, Sp4 and CmeA were added to *E. coli* W3110 MAGIC v.2. Both strains, *E. coli* W3110 Sp4 bioconjugation and *E. coli* W3110 SP4 MAGIC v.2, designated as gCmeA bioconjugation Sp4, and gCmeA MAGIC v.2 Sp4, respectively for ease, were grown overnight in media with shaking, and subcultured the following day. Biological conjugation was initiated by induction of *pglB* expression at OD_600_ of O.4-0.5 by 1 mM IPTG. CmeA 6xHis was purified via IMAC, then analyzed by Western blot analysis. CmeA from Sp4 bioconjugation and MAGIC v.2 Sp4, reacted positively when probed with Sp4 antibody. Western blot analysis showed a distinctive ladder indicative of glycosylation of CmeA by Sp4. Length variability of the glycan polymer was clearly noticed, where gCmeA Sp4 MAGIC v. 2 exhibits a longer polymer when compared to gCmeA bioconjugation Sp4 Fig 1, E and Fig S2. Image analysis of three biological replicates shows that the CmeA glycoconjugate increased by 3-fold in MAGIC v.2 when compared to traditional bioconjugation method, with glycosylation efficiency increased from 81.2 % ± 7 to 90.4 % ± 2.9. This result confirms that glycoprotein production is significantly enhanced by the MAGIC platform Fig 1, F.

We noticed that the efficiency of *E. coli* MAGIC strains construction was dependent on minimising the size of the cargo within the MAGIC constructs. In order to refine the MAGIC technology, we opted to assemble an OST under control of a constitutive promoter instead of an IPTG inducible promoter. We synthetically designed MAGIC v.3 utilizing the σ^70^ promoter(s) from Registry of Standard Biological Parts and the iGEM inventory number BBa_J23109 (0.04), BBa_J23114 (0.1), BBa_J23115 (0.15) and BBa_J23104 (0.72), where promoter strength was previously measured in the relative fluorescence units from plasmids expressing RFP in strain TG1 grown in LB media to saturation (23). Four promoters were chosen to cover a wide range of strength to drive the expression of the OST. Additionally, the inducible promoters assisted in further reducing the distance between the I and O end of the transposon by removing the P_tac_ and *lacI* repressor, potentially further reducing vaccine production cost by omitting the use of a protein induction reagent such as IPTG. Next, we assessed the impact of promoter strength on glycoconjugate production yield. We constructed *E. coli* CLM24 MAGIC v.3 variants and tested their relative ability to produce a vaccine against our model O-antigen; *F. tularensis* using CmeA as a carrier protein. Surprisingly, strong promoter BBa_J23100 (0.72) had a deleterious effect on bacterial growth and reduced the glycoconjugate yield when compared to the other three promoters. No growth defects were observed among the other *E. coli* MAGIC v.3 variants (BBa_J23109 (0.04), BBa_J23114 (0.1), BBa_J23115 (0.15) upon overnight growth at shaking media. Cells were subcultured the following day and proceeded to IMAC. Western blot analysis showed a higher glycoconjugate yield in CmeA isolated from *E. coli* MAGIC v.3 BBa_J23115 (0.15) (denoted as gCmeA PglB 0.15) when compared to the other promoters and *E. coli* CLM24 CmeA bioconjugation as a control (denoted as gCmeA PglB (0.04), gCmeA PglB (0.1), and gCmeA bioconjugation) Fig 1, G. The increase in glycoconjugate yield was reproducible in three biological replicates. Glycoprotein yield was estimated using image densitometry (glycoprotein/glycoprotein + unglycosylated protein *100) from three biological replicates Fig S3. When compared to the three plasmid bioconjugation method, gCmeA PglB (0.04) showed minimal glycosylation of CmeA. Interestingly, gCmeA PglB (0.1) showed a 1.6-fold increase in glycoprotein yield whilst gCmeA PglB (0.15) showed a 2-fold increase, when both compared to CmeA bioconjugation Fig 1, H. Glycosylation efficiency of CmeA were 71.9% ± 0.7 in gCmeA bioconjugation method, 74.8% ± 0.9 in gCmeA PglB (0.1), and 74.4% ± 0.6 in gCmeA PglB (0.15) These results demonstrate the importance of fine-tuning glycoengineering components to achieve optimum glycoconjugate yield. Taken together these results demonstrate that MAGIC rapid and robust method to advance glycoconjugate production in different glycoengineering *E. coli* strains.

### Developing of a candidate conjugate vaccine against *E. coli* O157

One of the most challenging steps in bioconjugation and cell free glycosylation methods is the successful expression of the glycan orthogonal pathway in *E. coli*. This problem is further complicated when ORFs of a certain glycan are scattered on the genome and/or when a certain degree of acetylation is necessary for a carbohydrate to be immunogenic (24) and/or the acetyltransferase responsible for this is unknown. In order to overcome this bottleneck, we evaluated the robustness of the MAGIC platform in the development of glycoconjugates in a non-*E. coli* strain. The bacterium *C. sedlakii* NRC6070 is a non-pathogenic bacterium that expresses an identical O-antigen to enterohemorrhagic *E. coli* O157, a food-borne pathogen that causes haemolytic-uremic syndrome (HUS) and haemorrhagic colitis, with infectious dose as low as 10^2^ CFU (25,26). Antibiotic treatment of *E. coli* O157 could increase the potential risk of development HUS. Phase II .clinical trials showed that an O157:H7 O-antigen conjugated to exotoxin A from *Pseudomonas aeruginosa* was safe and immunogenic, as it induced serum IgG LPS antibodies approximately 20-fold higher than pre-vaccination titres after 26 weeks post immunization(25). The O-antigen consists of the tetrasaccharide repeating unit; α-PerNAc-(1→3)-α-Fuc-(1→4)-β-Glc-(1→3)-α-GlcNAc (26).

To assess the versatility and robustness of MAGIC in developing a candidate vaccine in the event of a potential outbreak situation, we aimed at developing a candidate vaccine against foodborne pathogen *E. coli* O157:H7, as an exemplar. We took into consideration two main factors, firstly, to test a rapid method for vaccine development; secondly, to avoid growing a large culture volume of a pathogenic organism which can be a major safety biohazard (27). Since *E. coli* O157:H7 is categorized as a CAT III organism, we used *Citrobacter sedlakii* instead. We developed *C. sedlakii* MAGIC v.1 as described in Fig 2, A. This time we added a single plasmid expressing our model carrier protein, CmeA 6xHis, as *C. sedlakii* expresses the same O-antigen as *E. coli* O157:H7. The development of *C. sedlakii* MAGIC v.1 expressing CmeA took 3 days in total from streaking the cells in glycerol stocks. To achieve the second aspect, we sought to grow *C. sedlakii* MAGIC v.1 CmeA on 2 LB agar plates supplemented with 5 µM IPTG. The following day, cells were scraped, washed twice with PBS, and CmeA was IMAC purified. Western blot analysis of CmeA isolated from *C. sedlakii* MAGIC v.1 showed an increase in the molecular weight of CmeA in the form of a clear double bands that were not seen in CmeA isolated from *C. sedlakii* wildtype Fig 2, B. The increase in mass shift is generally seen as an indication of glycosylation. To confirm this finding, we probed CmeA variants with O157 monoclonal antibody (abcam), this identified a high molecular weight ladder reacting positively in CmeA isolated from *C. sedlakii* MAGIC v.2 but not in the CmeA *C. sedlaki* wildtype. Interestingly, the double bands did not react with the monoclonal antibody Fig 2, C.

**Figure 2.**
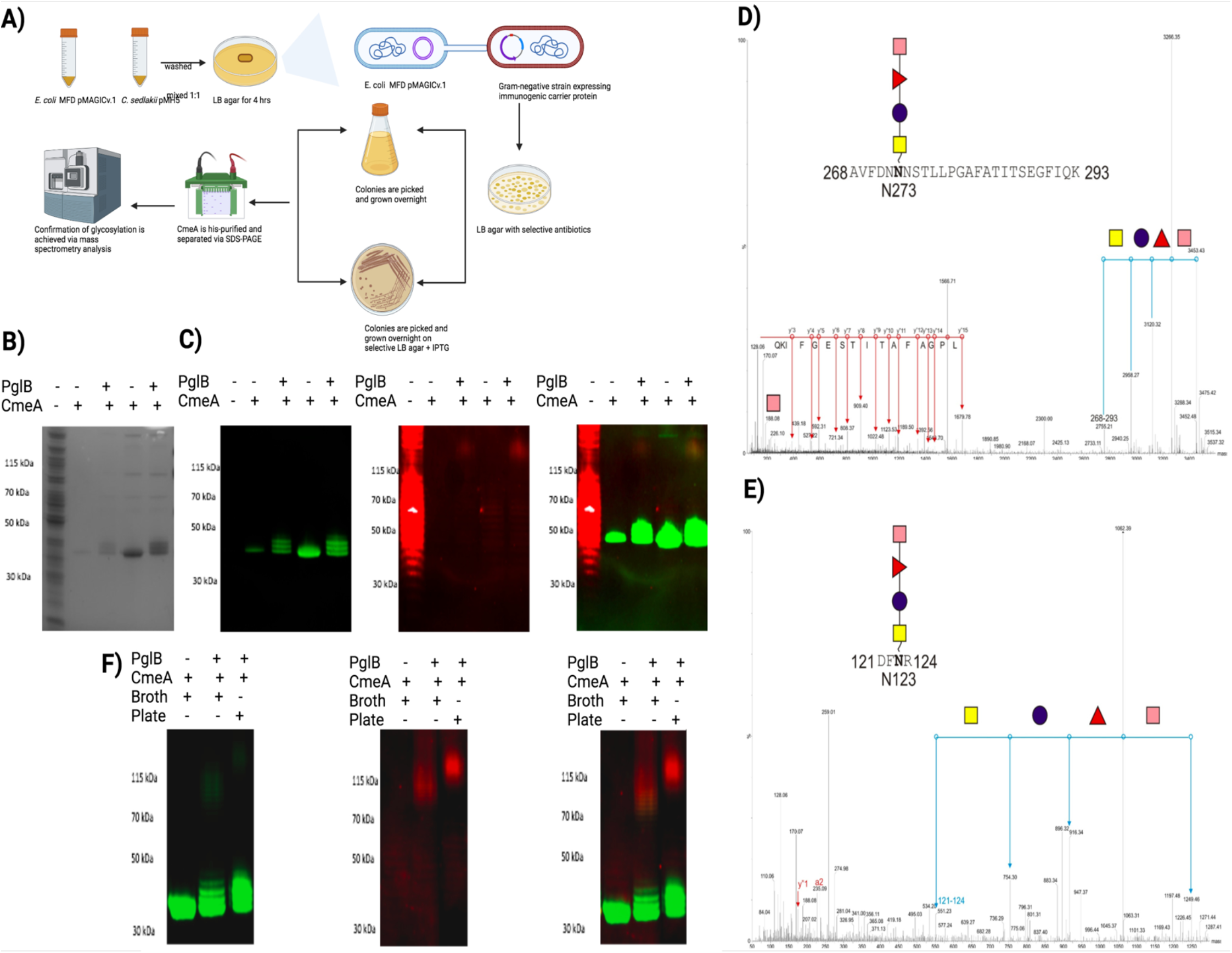
Developing of *E. coli* O157 candidate conjugate in *C. sedlakii* MAGIC v.1. A, schematic diagram of construction of *C. sedlakii* MAGIC v.1; B, Coomassie stain of His-tagged CmeA protein purified from *C*.*sedlakii* and *C. sedlakii* MAGIC v.1 by nickel affinity chromatography. Biological samples were separated on a Bolt 4-12% bis-tris gel (Invitrogen) with MOPS buffer; C, western blot analysis of CmeA purified from *C. sedlakii* and *C. sedlakii* MAGIC v.1, Biological samples were separated on a Bolt 4-12% bis-tris gel (Invitrogen) with MOPS buffer transferred to nitrocellulose membrane with an iBlot 2 dry blotting system. The membrane was probed with anti-His (Invitrogen) and anti-O157 (abcam) antibody and detected with fluorescently labelled secondary antisera (green-His, red-O157) on a LiCor Oddysey scanner; D, Mass spectrometry analysis of CmeA glycopeptides ^268^AVFDN**N**NSTLLPGAFATITSEGFIQK^293^; and E^121^DF**N**R^124^ ; purified CmeA was reduced, alkanylted, and treated with sequencing grade trypsin overnight, peptides were then run on LC-MS/MS (Waters). Precursor ion fragmentation shows the loss of HexNAc, Hex, deoxyHex, and deoxyHexNAc which corresponds to one repeating unit of O157. Glycan fragmentation is shown in blue lines and peptide fragmentation is shown in red; F, glycosylation of CmeA in *C. sedlakii* MAGIC v.1 in broth and plate of His-tagged CmeA protein purified from by nickel affinity chromatography. Biological samples were separated on a Bolt 4-12% bis-tris gel (Invitrogen) with MOPS buffer; C, western blot analysis of CmeA purified from *C. sedlakii* and *C. sedlakii* MAGIC v.1, Biological samples were separated on a Bolt 4-12% bis-tris gel (Invitrogen) with MOPS buffer transferred to nitrocellulose membrane with an iBlot 2 dry blotting system. The membrane was probed with anti-His (Invitrogen) and anti-O157 (abcam) antibody and detected with fluorescently labelled secondary antisera (green-His, red-O157) on a LiCor Oddysey scanner.

We used LC-MS/MS analysis to precisely characterize the double bands observed in CmeA *C. seldakii* MAGIC v.2. Gel slices were cut followed by reduction, alkylation, and digested with trypsin. Peptides were separated by LC and detected by CID MS/MS. CmeA was identified after the raw data search. Further data analysis was performed with the addition of O157:H7 repeating unit molecular weight (698 Da) to modification list in the database search method. Two peptides were identified as a part of the D/E-X-N-X-S/T (where X is any amino acid except proline) carrying this glycan ^268^AVFDN**N**NSTLLPGAFATITSEGFIQK^293^; *m/z* 3452 and ^121^DF**N**R^124^ ;*m/z* 1249). Manual analysis of the MS/MS data showed fragmentation of the peptides with characteristic peak (*m/z* 188) in both peptides, which is indicative of PerNAc loss. Further analysis of the peaks identified the modification by a tetramer of 187-146-162-203. The mass of 187 is consistent with an *O-*acetyl deoxyhexose, 146 is consistent with deoxyhexose, 162 is consistent with a hexose, and 203 is consistent with *N-*acetyl hexosamine Fig 2, D-E. These data combined with the Western blot analysis confirm that CmeA is successfully glycosylated with the correct O157 sugar residue by MAGIC v.2.

Next, we sought to assess if there is a difference in glycoconjugate phenotype between an agar plate glycosylation method and the traditional liquid culture media method. Cells were grown, and glycosylation was induced as mentioned in the methods section CmeA 6xHis was affinity purified and analyzed by western blot. CmeA from both culture media and plate showed the distinct extra high molecular weight bands indicative of glycosylation with O157-Ag. Interestingly, in liquid culture media conditions, CmeA exhibited lower polymer length compared to plate glycosylation method. Both of the polymers reacted positively when probed with O157 monoclonal antibody Fig 2, F. These results clearly show that MAGIC is a robust and versatile method that could be used in the development of candidate vaccines against bacterial pathogens in a non-*E. coli* strain. Additionally, we demonstrate that the biotechnology is compatible with health and safety procedures that minimize any biohazard risk.

## DISCUSSION

In this work we have established MAGIC, as a versatile and robust, “plug-and-play” platform for vaccine development. Using *F. tularensis* CmeA-Ft-OAg as a model, we improved the bacterial growth, resulting in a higher biomass and increased glycoconjugate(s) yield. The enhancement of glycoconjugate production was evident in degree of polymerization visible in *F. tularensis* conjugate vaccine developed using the MAGIC integrated strain. This degree of polymerization was not achieved either by traditional bioconjugation or cell free glycosylation methods Fig S4. Degree of polymerization is one of the factors that could impact vaccines efficacy. In a recent study, large molecular size Ft-OAg polysaccharide glycoconjugate provided superior protection than low molecular size glycoconjugate against intranasal *F. tularensis* challenge in a mouse model of tularemia (28). Previous attempts to produce highly polymerized glycoconjugates using bioconjugation methods were hampered by expressing several glycoengineering components in the cell from plasmids, which consequently, led to lower cellular biomass and glycoconjugate yield. By alleviating the metabolic stress in the cell when applying MAGIC to one or more glycoengineering components, cellular biomass could increase leading to a higher glycoconjugate yield. Indeed, cellular biomass increased by 41% when MAGIC was applied, compared to traditional bioconjugation three plasmids method. This increase in biomass was accompanied by improved glycosylation efficiency, seen as an increase in the polymer length in the glycoconjugate, and a significant increase in the glycoconjugate yield.

In-depth analysis of the vaccine market shows that pneumococcal conjugate vaccine (PCV) is the most likely to see a high value growth by 2030, requiring billions of doses to be manufactured (2). The instrumental role played by MAGIC in developing another efficacious and inexpensive vaccine is exemplified by the *S. pneumoniae* glycoconjugate generated. When compared to the three plasmid bioconjugation method, the MAGIC assembled strain showed a significant 3-fold increase in glycoconjugate yield with glycosylation efficiency reaching 90.4 % ± 2.9. Additionally, glycoconjugates produced using the MAGIC platform showed a higher degree of polymerization of SP4 glycan when compared to traditional bioconjugation method. When tested in outbred mice these glycoconjugates had significant immunogenicity and protection resulting in higher survival rate than animals immunized with PCV13 (low dose) following challenge with *S. pneumoniae* (14). Taken together, we set out a novel benchmark in biological conjugation that allows for enhancement of glycoconjugates production with key advantages over the current biological conjugation technologies.

In contrast to the recently published cell free glycosylation method, MAGIC can provide an inexhaustible and renewable source of glycoconjugates (29). This main difference stems from the fact that MAGIC is based on converting the bacterial cell, either *E. coli* or any other Gram-negative bacterium, into a factory for glycoconjugate vaccine production, contrary to the limited reaction volumes in cell free glycosylation (30). One appealing feature of bioconjugation is the reduction of the batch-to-batch variation bottleneck, since no component mixing is required with specific quantities that increase the probability of a human error (29). Key steps are also eliminated when compared to cell free methods such as, ultracentrifugation, protein and LLO quantification (30).

The need to develop vaccines in an outbreak situation has been dramatically demonstrated on a global scale through the development of vaccines against COVID-19 using mRNA and adenovirus-based technologies. These technology platforms are less suitable for tackling most bacterial infections. Therefore, we devised a hypothetical bacterial outbreak scenario, and demonstrated that MAGIC could be an indispensable tool in developing a candidate vaccine in short time. We demonstrated for the first time that glycosylation could be achieved by growing cells in plates. This eliminates the need to grow large cultures of pathogens for chemical conjugation and cell free glycosylation. Additionally, it prevents potential health risks associated with culture spillage which could cause a disease outbreak. Our outbreak scenario of developing a candidate vaccine against *E. coli* O157 was achieved in one week starting from streaking the wildtype strain(s) to purifying a glycoconjugate. Such acceleration in developing of glycoconjugate highlights the biotechnological potential of the MAGIC platform and its general applicability.

According to the World Health Organization antibiotic resistant bacteria cause around 700,000 deaths annually and has been called the slow pandemic. By 2050, this number is expected to be 10 million (31). There is a global imperative to develop biotechnology platforms for cheap and effective vaccines against antibiotic resistant bacteria. The advances in chemoenzymatic conjugation, synthetic glycobiology, and bacterial glycosylation have led to breakthroughs in vaccine development. To this end, we designed MAGIC to allow the testing of different bioconjugation components and/or different glycosylation systems, simultaneously, in a quick and efficient way. We anticipate that MAGIC could solve several obstacles in vaccine scalability such as, the choice of plasmids with compatible origin of replications, avoiding antibiotic usage, elimination of any induction chemical, thus reducing the total cost of vaccine production. Metabolic studies and glycomics will lead the fine-tuning of MAGIC technology in glycoengineering *E. coli*. This will help to glyco-customise bacterial strains for specific purposes; not only to test a repertoire of OSTs for diverse glycan structures and linkages, but also different carrier protein combinations.

In summary, we present a novel glycoengineering platform that will accelerate developing a range of glycoconjugate vaccines. The platform provides unique features such as, a) enhancing bacterial growth rate by decreasing the metabolic burden exerted by glycosylation components on the cell, b) glycoconjugate yield gains c) the ability to rapidly generate rationally designed tailormade vaccines, d) built-in modularity and compatibility with any glycosylation machinery available, and e) wide applicability in non-glycoengineering *E. coli* cells. Collectively, the application of MAGIC technology could be used in the improvement of existing vaccines, the development of new vaccines, and potentially as a rapid response vaccinology strategy in an outbreak situation.

## Acknowledgements

We would like to thank Steven Lynham, mass spectrometry centre of excellence, King’s college London, UK for his help with mass spectrometry. We thank the Biotechnology and Biological Sciences Research Council, including grants BB/M01925X/1, BBSRC BB/H017437/1 for helping to fund this research.

## FIGURES

**Figure S1.**
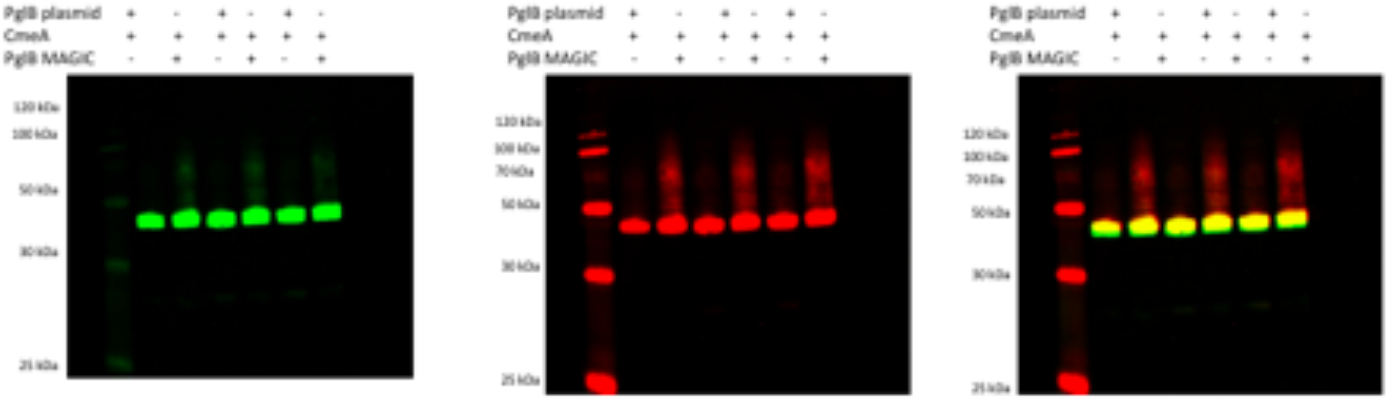
Western blot of 5 µg His-tagged CmeA protein purified by nickel affinity chromatography. Triplicate biological samples were separated on a Bolt 4-12% bis-tris gel (Invitrogen) with MOPS buffer and transferred to nitrocellulose membrane with an iBlot 2 dry blotting system. The membrane was probed with anti-His (Invitrogen) and anti-Ft-O antigen monoclonal antibody (abcam) and detected with fluorescently labelled secondary antisera (green-His, red-Ft-O-antigen) on a LiCor Oddysey scanner.

**Figure S2.**
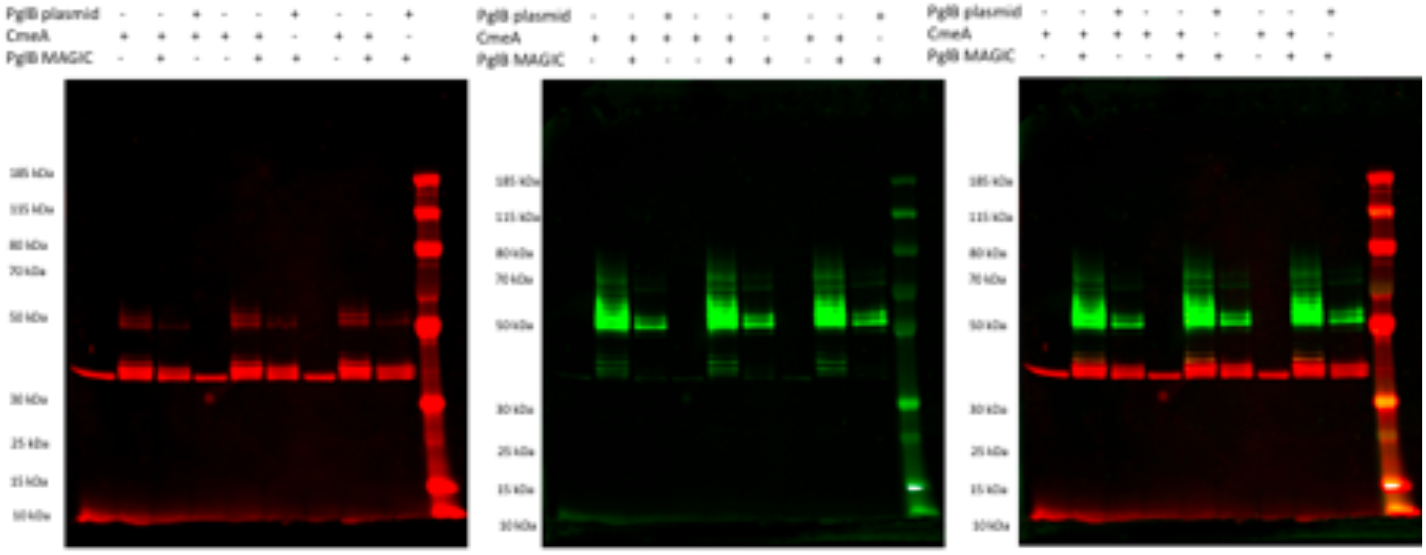
Western blot of 5 µg His-tagged CmeA protein purified by nickel affinity chromatography. Triplicate biological samples were separated on a Bolt 4-12% bis-tris gel (Invitrogen) with MOPS buffer and transferred to nitrocellulose membrane with an iBlot 2 dry blotting system. The membrane was probed with anti-His (Invitrogen) and anti-SP4 (Statens) antisera and detected with fluorescently labelled secondary antisera (red-His, green-SP4) on a LiCor Oddysey scanner.

**Figure S3.**
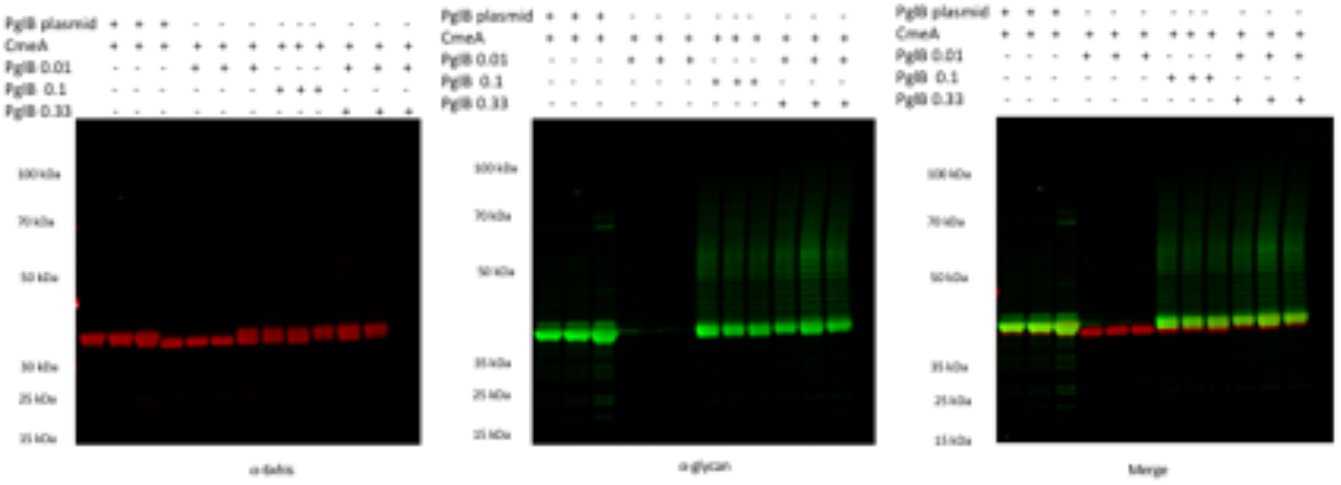
Western blot of 5 µg His-tagged CmeA protein purified by nickel affinity chromatography. Triplicate biological samples were separated on a Bolt 4-12% bis-tris gel (Invitrogen) with MOPS buffer and transferred to nitrocellulose membrane with an iBlot 2 dry blotting system. The membrane was probed with anti-His (Invitrogen) and anti-Ft-O antigen monoclonal antibody (abcam) and detected with fluorescently labelled secondary antisera (red-His, green-Ft-O-antigen) on a LiCor Oddysey scanner.

**Figure S4.**
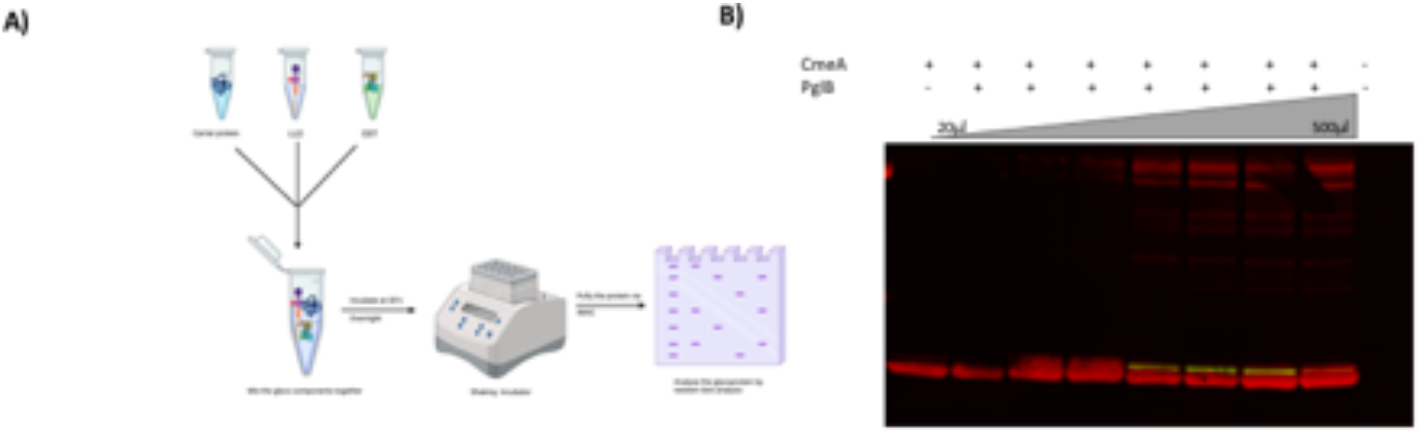
Cell free glycosylation (CFG). A, schematic diagram of cell free glycosylation analysis; B, western blot of CFG, Western blot of 5 µg His-tagged CmeA protein purified by nickel affinity chromatography. Triplicate biological samples were separated on a Bolt 4-12% bis-tris gel (Invitrogen) with MOPS buffer and transferred to nitrocellulose membrane with an iBlot 2 dry blotting system. The membrane was probed with anti-His (Invitrogen) and anti-Ft-O antigen monoclonal antibody (abcam) and detected with fluorescently labelled secondary antisera (red-His, green-Ft-O-antigen) on a LiCor Oddysey scanner.

## METHODS

### MAGIC construction

The gene coding for *C. jejuni* NCTC11168 PglB was loaded into a Mini-Tn*5*Km2 transposon within a pUT backbone targeting the NotI site. However, in order to assemble a construct that could be induced, we first cloned *C. jejuni pglB* into pEXT20. This is a vector that enables IPTG inducible expression of ORFs that are inserted within the MCS ^14^. The gene coding for *C. jejuni* PglB was amplified by PCR from the plasmid pACYC*pgl* with the pTac promoter and LacIq repressor from plasmid pEXT20 as a template using Pfx Polymerase (Thermo Fisher Scientific UK) with (SEQ ID 15: 5′-TTTTGCGGCCGCTTCTACGTGTTCCGCTTCC-3′) as forward primer and (SEQ ID 16: 5′-TTTTGCGGCCGCATTGCGTTGCGCTCACTGC-3′) reverse primer using the following cycling conditions, 94 °C 2 min followed by 35 cycles of 94 °C for 30 sec, 56 °C for 30 sec and 68 °C for 4 min. The PCR product was then cloned in pJET2.0 plasmid (Thermo Scientific U.K.) according to the manufacturer’s instructions and named pOST9. The plasmid was maintained in *E. coli* DH5α cells (Stratagene U.K.). The vector pOST9 was cut with the restriction enzyme NotI (New England Biolabs U.K. Ltd.) and ligated into the unique NotI site in pUTMini-Tn*5*Km2 resulting in plasmid pJAN25 and maintained in Transformax *E. coli* strain EC100D *pir*+ (Cambio U.K.). The plasmid was then transformed into *E. coli* 19851 *pir*+ or *E. coli* MFD for maintenance and for conjugation with host bacteria for delivery on *pglB*.

### Bacterial Conjugation

To enable transfer of the *CjpglB* and *cmeA* from the transposon cargo into the chromosome of a recipient *E. coli* strain using the plasmids pJAN25 and pFEB11 respectively, the new loaded Mini-Tn*5*Km2 transposons were maintained in *E. coli* strain19851*pir*^+^. We switched to using *E. coli* MFD a diaminopimelic acid (DAP) auxotroph. Growth medium was supplemented with kanamycin 50 µg/ml and ampicillin 100 µg/ml for pJAN25 or pFEB11 whilst chloramphenicol 30 µg/ml and ampicillin 100 µg/ml were added to maintain pEFNOV19. Both donor and recipient bacteria were growth until late exponential phase. Bacterial cells were pelleted by centrifugation, washed 3 times with PBS and mixed together in a ratio of 1:3 recipient to donor and spotted on a dry LB agar plate with no antibiotics for 4-8 hrs. The cells were scraped and suspended in PBS and dilutions plated on LB agar with appropriate selection antibiotics to select for transconjugants. Individual colonies were picked up and screened for loss of the pUT backbone and for the presence of the transposon.

### Protein purification

Protein purification was carried out using QIAExpressionist NiNTA purification kit according manufacturer’s instructions (Qiagen, Germany) and following the HIS purification under native conditions protocol. Purification was carried out from 10 ml of bacterial culture. If induced, bacterial cells from an o/n culture were used to inoculate 10 ml of Luria Bertani (LB) broth. Cells were incubated at 37 °C, 180 rpm until an OD_600_ of 0.4 was reached. At this point L-arabinose was added to a 0.2% v/v final concentration or IPTG at a 1mM final concentration. Samples were incubated at 37 °C for a further 16 h before the cell pellet was collected for protein purification. Cells were lysed in lysis buffer (50 mM NaH_2_PO4, 300 mM NaCl, 10 mM Imidazole, pH 8) following manufacturers protocol. Cells debris were removed by centrifugation at 12,000 xg, supernatant was incubated for an hour with NiNTA agarose at 4 °C then 4 times beads volume with wash buffer (50 mM NaH_2_PO4, 300 mM NaCl, 20 mM Imidazole, pH 8), then eluted with 50 mM NaH_2_PO4, 300 mM NaCl, 250 mM Imidazole, pH 8.

### Mass spectrometry

In gel reduction, alkylation, and digestion with trypsin or chymotrypsin was performed on the gel sample prior to subsequent analysis by mass spectrometry. Cysteine residues were reduced with dithiothreitol and derivatized by treatment with iodoacetamide to form stable carbamidomethyl derivatives. Trypsin digestion was carried out overnight at room temperature after initial incubation at 37°C for 2 h. Sample digests, resuspended in 0.1% (v/v) formic acid, were analyzed by on-line nano-flow reverse-phase high-performance liquid chromatography with online electrospray-mass spectrometric analysis (nano-RP-HPLC-ES-MS) with MS/MS (MS^e^) using a Waters SYNAPT G2-S high-definition mass spectrometer, coupled to a Waters ACQUITY UPLC M-Class System (Waters UK, Elstree). Separations were achieved by means of a C18 trapping column (M-Class Symmetry C18 Trap, 100 Å, 5 µm, 180 µm × 20 mm, 2G) connected in-line with a 75 µm C18 reverse-phase analytical column (M-Class Peptide BEH C18, 130 Å, 1.7 µm, 75 µm × 150 mm) eluted over 90 min with a gradient of acetonitrile in 0.1% formic acid at a flow rate of 300 nL/min. Column temperatures were maintained at 50°C, and data were recorded in MS^e^ “Resolution” positive ion mode, with scan times set to 0.5 s in both the high-energy and low-energy modes of operation. The instrument was pre-calibrated using 10–100 fmol/µL of [Glu^1^]-fibrinopeptide B/5% (v/v) acetic acid (1:3, v/v) and calibrated during analysis by means of a lockmass system using [Glu^1^]-Fibrinopeptide B 785.8426^2+^ ion. The collision gas utilized was argon with collision energy ramp of 20–45 eV. Data acquisition was performed using MassLynx (Waters UK, Elstree) software and analyzed by means of MassLynx, BiopharmaLynx and ProteinLynx Global Server (PLGS) version 3.0.2 (Waters UK, Elstree).

### Cell free glycosylation

Cell free glycosylation was conducted in S30 buffer with 0.1% n-dodecyl-β-d-maltopyranoside (DDM; Thermo Scientific) and 10 mM MnCl_2_ (Across Organics). A constant volume of lysed acceptor protein and glycan was mixed to varying volumes of lysed OSTs ranging from 100 µl to 500 µl for a total reaction volume of 1 ml. Cell free glycosylation was conducted at 30 °C and 110 rpm overnight. Afterwards, the samples were centrifuged at 12,000 xg for 10 minutes to remove reaction debris. The samples were then incubated for an hour and a half in Ni-NTA resin (Qiagen) at 4 °C to pull down glycosylated acceptor protein. After washing samples with His Wash buffer (50 mM NaH_2_PO4, 300 mM NaCl, 20 mM Imidazole, pH 8), the samples were eluted using 75 µl His Elution buffer (50 mM NaH_2_PO4, 300 mM NaCl, 250 mM Imidazole, pH 8).

## Supplementary method

### Assembly of constitutively expressed *C. jejuni* PglB

Using *C. jejuni* 81116 genomic DNA as a template pglB was amplified using the forward primers

X4

5’- TTTTGCGGCCGCTTTACAGCTAGCTCAGTCCTAGGGACTGTGCTAGCAGGAGGAAAAAAATGTTGAAAAAAGAGTATTTAAAAA -3’ BBa_J23109

X10

5’- TTTTGCGGCCGCTTTATGGCTAGCTCAGTCCTAGGTACAATGCT AGCAGGAGGAAAAAAATGTTGAAAAAAGAGTATTTAAAAA -3’ BBa_J23114

X15

5’- TTTTGCGGCCGCTTTATAGCTAGCTCAGCCCTTGGTACAATGCTAGCAGGAGGAAAAAAATGTTGAAAAAAGAGTATTTAAAAA -3’ BBa_J23115

X72

5’-TTTTGCGGCCGC TTGACAGCTAGCTCAGTCCTAGGTATTGTGCTAGCAGGAGGAAAAAAATGTTGAAAAAAGAGTATTTAAAAA-3’ BBa_J23104

REV *CjpglB*: 5’- TTTTGCGGCCGCTTAAATTTTAAGTTTAAAAACTTTAGC-3’

94°C/15 s, [94 °C/30 s, 50 °C/30s, 68 °C/2min]* 68°C/2min

*35 cycles

PCR products were digested with NotI HF (New England Biolabs) and cloned into NotI digested Cloned into pUC57ZeoTn.

The plasmid was then digested with SfiI and ligated to pUTminiTn*5* to create the plasmid pELLA1 (pglBx4, pELLA2 (pglBx10), pELLA3 (pglBx15). Conjugation to the *E. coli* strain CROW was carried out to deliver the constitutive *pglB* copy.

**Table 1.** Strains and plasmids used in this study

Strains and plasmids used in this study
| Strains/Plasmids | Description | Reference or source |
| --- | --- | --- |
| *E. coli* DH5α | F- ϕ80*lacZ*ΔM15 $\Delta$(*lacZYA-argF*) U169 *deoRrecA1 endA1 hsdR17* (rk-, mk+) *gal-phoA* supE44 λ-*thi*-1 *gyrA96 relA1* | Thermoscientific (UK) |
| *E. coli* W3110 | *rph-lIN(rrnD-rrnE)* | (19) |
| *E. coli* CLM24 | *rph-lIN(rrnD-rrnE)* 1 Δ*waaL* | (19) |
| *E. coli* SΦ874 | $\Delta$(*sbc-rfb*)86 O75<sup>-</sup> | (32) |
| *E. coli* SCM3 | SΦ874 Δ*waaL* | (33) |
| *E. coli* SCM6 | SΦ874 Δ*waaL* Δ*wecA* | Miguel Valvano (Queen's University, Belfast, UK) |
| *E. coli* SCM7 | SΦ874 Δ*wecA* | Miguel Valvano (Queen's University, Belfast, UK) |
| *E. coli* SDB1 |  | (12) |
| *Citrobacter sedlakii* | prototroph | ATCC |
| *E. coli* CROW | *rph-lIN(rrnD-rrnE)* 1 Δ*waaL* , Δ*lpxM* | Kay, E *et al* submitted |
| *E. coli* W3110 MAGIC v.1 | *E. coli* W3110 P<sub>tac</sub>*pglB* kan<sup>r</sup> | This study |
| *E. coli* CLM24 MAGIC v.1 | *E. coli* CLM24 P<sub>tac</sub>*pglB* kan<sup>r</sup> | This study |
| *E. coli* SΦ874 MAGIC v.1 | *E. coli* SΦ874 P<sub>tac</sub>*pglB* kan<sup>r</sup> | This study |
| *E. coli* SCM3 MAGIC v.1 | *E. coli* SCM3 P<sub>tac</sub>*pglB* kan<sup>r</sup> | This study |
| *E. coli* SCM6 MAGIC v.1 | *E. coli* SCM6 P<sub>tac</sub>*pglB* kan<sup>r</sup> | This study |
| *E. coli* SCM7 MAGIC v.1 | *E. coli* SCM7 P<sub>tac</sub>*pglB* kan<sup>r</sup> | This study |
| *E. coli* SDB1 MAGIC v.1 | *E. coli* SDB1 P<sub>tac</sub>*pglB* kan<sup>r</sup> | This study |
| *Citrobacter sedlakii* MAGIC v.1 | *C. sedlakii* P<sub>tac</sub>*pglB* kan<sup>r</sup> | This study |
| *E. coli* CROW MAGIC v.3 | (*pglB* is controlled by BBa_ BBa_J23109<br>BBa_J23114, BBa_J23115) | This study |
| *E. coli* W3110 MAGIC v.2 | *E. coli* W3110 P<sub>tac</sub>*pglB* | This study |
| pUT-mini-Tn5Km2 | *ori* R6K, mob RP4, miniTn5km-based delivery with delivery plasmid with Amp<sup>r</sup> | (34) |
| pGAB2 | *Francisella tularensis* O-antigen biosynthesis locus cloned in pLAFR-1 | (10) |
| pB4 | *S. pneumoniae* serotype 4 capsule locus (*wciI-fnlC*) pBBR1MCS-3 (Tc<sup>R</sup>) | (14) |
| pWA2 | *C. jejuni* CmeA with a pelB signal sequence and His6-tag. (Ap<sup>R</sup>) | (35) |
| pJAN25 | *ori* R6K, mob RP4, miniTn5*km*-based delivery with delivery plasmid with Amp<sup>r</sup> coding for IPTG inducible *C. jejuni* pglB cloned in NotI site controlled by pTac promoter. | This study |
| pUA73 | pTac promoter. Zeor cassette flanked by loxP sites | This study |
| pOST9 | pMB1 *ori*, Amp<sup>r</sup> plasmid carrying IPTG inducible *C. jejuni* pglB | This study |
| pELLA1 | *ori* R6K, mob RP4, miniTn5*Zeo*-based delivery with delivery plasmid with Amp<sup>r</sup> coding for IPTG inducible *C. jejuni* pglB cloned in NotI site controlled by constitutive promoter BBa_J23109 | This study |
| pELLA2 | *ori* R6K, mob RP4, miniTn5*Zeo*-based delivery with delivery plasmid with Amp<sup>r</sup> coding for IPTG inducible *C. jejuni* pglB cloned in NotI site controlled by constitutive promoter BBa_J23114 | This study |
| pELLA3 | *ori* R6K, mob RP4, miniTn5*Zeo*-based delivery with delivery plasmid with Amp<sup>r</sup> coding for IPTG inducible *C. jejuni* pglB cloned in NotI site controlled by constitutive promoter BBa_J23115 | This study |

